# Design and structure of protein cages based on helical fusion and machine learning

**DOI:** 10.64898/2026.05.27.727936

**Authors:** Pablo San Segundo-Acosta, Johanne Le Coq, Jasminka Boskovic, Robin A. Aglietti, Peter Bowers, Todd O. Yeates, Roger Castells-Graells

## Abstract

Self-assembling protein cages are versatile nanoscale architectures with broad applications in drug delivery, vaccine development, and structural biology. Historically, two main strategies have been used to construct such cages: genetic fusion of oligomeric domains connected by helical linkers, and computational interface design using either physics-based or machine learning-based methods. Here, we extend the original fusion approach using modern AI algorithms and more sophisticated treatments of helix bending to create protein cages with novel architectures composed exclusively of trimeric building blocks arranged in tetrahedral symmetry. Of fifteen designs tested experimentally, multiple sequence variants of two of these designs assembled predominantly into soluble, monodisperse particles of the expected size, with native molecular masses of 633 kDa (T33-Fus-1A, B) and 638 kDa (T33-Fus-2). Cryo-electron microscopy (cryo-EM) structures of three distinct sequence variants spanning from 3.0-3.9 Å in resolution confirmed the intended structures in atomic detail, with C-alpha RSMD values over the entire assemblies as low as 2 Å. The predicted modes of helix bending were similarly validated. The results highlight the impact of methodological improvements for achieving a level of regularity and design precision that has largely evaded prior applications of the fusion approach. These findings expand the prospects and accessible design space for self-assembling protein nanomaterials.

## INTRODUCTION

Protein cages are symmetric macromolecules that self-assemble from multiple copies of one or more protein subunits. Their unique architecture, typically combining a defined interior cavity with a programmable exterior surface, has motivated extensive efforts to design them for applications in biotechnology and medicine (1–13).

The design concepts and first successful strategy for creating novel protein cages were introduced by Padilla, Colovos, and Yeates (2001) (14), who recognized that fusing two oligomeric protein domains through a rigid α-helical linker could enforce specific geometric relationships between their symmetry axes. When two (or more) axes of symmetry are brought into orientations that intersect at certain angles, multiple copies of the resulting fusion protein are driven to self-assemble into a closed, cage-like particle with tetrahedral, octahedral, or icosahedral symmetry. The underlying geometric rules for diverse constructions have been laid out (14–16). Following the creation of a tetrahedral cage, the oligomeric fusion approach was subsequently extended to produce a cubic cage with octahedral symmetry (17). In both cases, and in related work by others (18–20), challenges related to linker flexibility and the formation of somewhat distorted geometries or alternative assembly states highlighted the need for more precise control over the fusion geometry and the sequence of the connecting segments..

A complementary approach pioneered by King *et al*. (2012) (21), relies on computational design of de novo protein-protein interfaces to drive assembly. Physics-based methods using the Rosetta software suite enabled the creation of tetrahedral (21, 22), octahedral (21), and icosahedral (23) cages from one- and two-component systems. Motivated by the need to improve experimental success rates for design, machine learning (ML) algorithms were subsequently incorporated into numerous interface design studies. The introduction of ProteinMPNN (24) for sequence design has markedly improved experimental outcomes for designed protein nanomaterials (25–27). More recently, RFdiffusion (28) and related generative methods have opened the scope of de novo backbone generation, enabling the creation of higher-order symmetric and pseudosymmetric assemblies (29–32).

While modern computational tools have helped advance many interface design-based studies, comparable advances for fusion-based assembly approaches have been more limited (33–35). Here, we expand and improve upon previous fusion-based protein cage design work through two main advances: the use of machine learning methods to optimize key regions of the fusion constructs generated, and an algebra-based consideration of natural alpha helix bending. We describe a strategy combining classical mechanics and machine learning algorithms to create protein cages from two trimeric building blocks arranged in tetrahedral symmetry, with multiple successful designs confirmed by high-resolution cryo-EM structure determination.

## RESULTS

### Computational Design Pipeline for Trimer-Trimer Helical Fusion Cages

Our design strategy combines geometric connection of trimeric building blocks with ML-based sequence optimization at the helical junction and any other regions where incidental contacts might occur between the two trimeric components (Fig. 1). A tetrahedral cage composed of two distinct trimeric components (termed a T33 architecture) comprises 4 copies of each trimer type, with different types occupying alternating vertices of a cube (Fig. 1). If the two trimer types are situated at roughly similar distances from the center of the assembly, the overall architecture tends to resemble a cube. If one trimer type is more distal compared to the other, the overall shape tends to resemble a tetrahedron. When the T33 architecture is based on a genetic fusion of the two component types, the cage or nanoparticle is thus assembled from 12 identical copies of a subunit bearing the two component types connected by a linker, which is typically a continuous α-helix that spans between the two domains.

**Figure 1.**
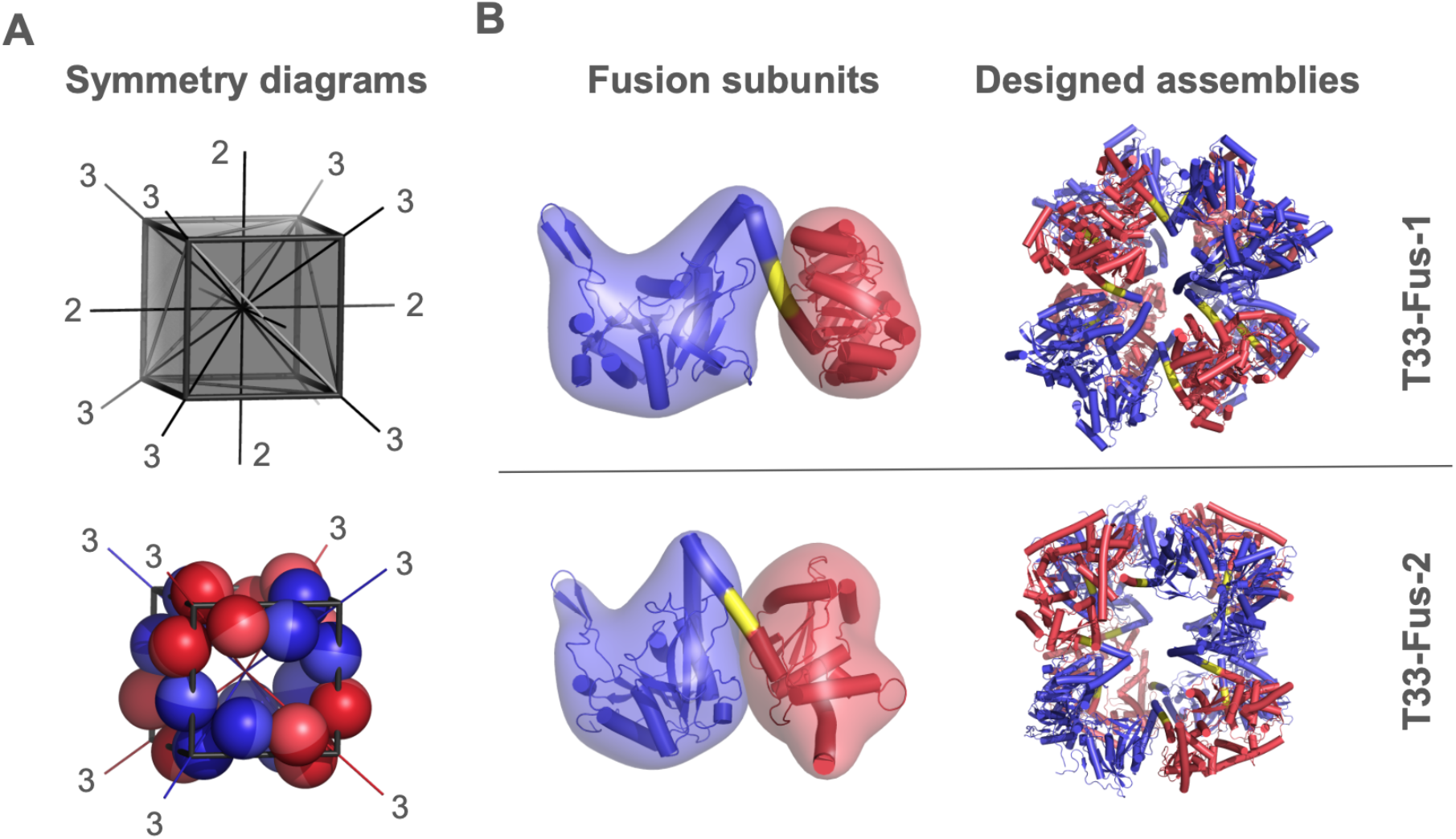
Design principles for trimer-trimer helical fusion cages and schematic models. **(A)** The symmetry axes for tetrahedral symmetry are shown on a cube (top) with two different types of trimeric components situated on alternating vertices (bottom). The 2-fold axes of symmetry, though they remain, are not drawn. **(B)** Elements of the design: two homotrimeric proteins of known structure with terminal α-helices are identified and connected in tetrahedral symmetry such that their helices point toward one another. A continuous helical linker bridges the two trimers, and the junction sequence is optimized using ProteinMPNN.

We began by mining the Protein Data Bank (PDB) (36) for homotrimeric proteins possessing terminal α-helices of at least 10 residues, reasoning that such helices provide the rigid structural element necessary for a predictable fusion geometry. Candidate trimers were required to have helices at either their N- or C-terminus. All pairwise combinations of a C-terminal-helical component and an N-terminal-helical component were then evaluated for consistency with tetrahedral point group symmetry after joining their ends by an alpha helical linker. The added linker was allowed to be a maximum of 7 residues to a minimum of −2; the latter case means the native helical termini are overlapping.

The underlying geometric requirement for tetrahedral symmetry is that the three-fold axes of symmetry carried by the two components must intersect in three-dimensional space, at an angle of precisely 70.5° (i.e.,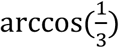. Thus, as a first step in design selection, combinatorial choices for the two components and the linker length were tested; the two termini were joined by an idealized alpha helix and the angle between the two symmetry axes thus generated were calculated (based on vector dot products) along with the closest approach distance between the two symmetry axes. Of course, the likelihood that any construct would satisfy such constraints exactly is zero, leading to an important challenge in design selection. In our initial fusion-based design studies (14, 37), we adopted a naive approach, essentially favoring design candidates where the two criteria (angle and separation distance between axes) were within some limits, with the expectation that a certain amount of helix bending would permit assembly. In the present study we employed a more structurally informed approach.

In our evaluation and selection of suitable candidates, we considered whether particular geometric perturbations – i.e., those needed to make a given design comply with the symmetry axis intersection requirements – could be accommodated by naturally allowable helix deformation. For illustration, consider a scenario where the geometric perturbation needed to meet the idealized angular design requirements – here, intersection at an angle of 70.5° – would require the connecting helix to be stretched along its axis. That mode of deformation is generally not observed in alpha helical segments, so even a minor deviation along that mode of deformation would not be a suitable starting point for design.

We developed an algebraic treatment of natural alpha helix variability to aid in evaluating design candidates. Briefly, we examined the variation across natural alpha helical structures using linear algebra and normal mode analysis. By characterizing the degree to which different modes of deformation are sampled in natural proteins, the presumptive helical deformation that would be required to idealize the symmetry axis requirements for any given design candidate could be evaluated against the allowable modes of alpha helix variability (see Methods). The resulting measure of helical deformation, as a pseudo energy, was used as the primary score for selecting designs.

Regarding other design metrics, candidate designs where the fusion geometry would lead to undue collisions between subunits were discarded. Cases where incidental contacts (or near-contacts) occurred were retained; in those cases, limited amino acid optimization was also undertaken at those positions. We further evaluated the interior cavity radius of candidate designs, discarding outlying cases with very open structures, which were judged to be subject to potentially problematic collapse. Finally, we restricted our attention to cases where both the N- and C-termini of the fusion subunit were exposed on the exterior surface, anticipating applications involving further fusion to additional peptides, antigens, or effector domains. Fifteen distinct design candidates were selected for sequence optimization by machine learning.

The ProteinMPNN model (24) was used to generate a series of sequence variants for each candidate design. Sequence design was performed in the context of intact 12-subunit symmetric assemblies. In all cases, the residues in the linking helix and any residues in the component domains in direct contact with those residues were subject to design. Any points of incidental contact between the multiple chains were also included in design. The remainder of the protein sequences were held fixed at their wild-type identities. For each design, 100 sequence designs were generated by ProteinMPNN, after which 2-6 constructs were chosen for experimental evaluation based on probability scores and similarity to the consensus sequence.

### Experimental Characterization of Designed Cages

Genes encoding the top 2-6 fusion protein designs across 15 sequence families were codon-optimized for expression in *Escherichia coli* and synthesized commercially. Proteins were expressed in BL21(DE3) LOBSTR cells and purified by immobilized metal affinity chromatography (IMAC) followed by size-exclusion chromatography (SEC). Of the 45 designs tested, five produced a prominent SEC peak eluting at a volume consistent with the expected molecular weight of the 12-subunit tetrahedral assembly, of which three were selected for follow up characterization. Hereafter those designs are referred to as T33-Fus-1A and T33-Fus-1B (~633 kDa with A or B referring to sequence variants) and T33-Fus-2 (~638 kDa). The T33-Fus-1 variants are based on fusion of PDB 3E35 and 4E38, T33-Fus-2 is based on fusion of PDB 2WAM and 3SYY. The remaining designs either failed to express in soluble form, eluted as aggregates in the void volume, or showed prominent peaks corresponding to monomeric or lower-order oligomeric species.

An analysis of amino acid sequences showed that the two sequence variants of T33-Fus-1 differ from each other at 6 positions, and from the starting fusion model (prior to ProteinMPNN) at 12 positions for T33-Fus-1A and 10 positions for T33-Fus-1B. For the second construct, T33-Fus-2 differs from the starting fusion model at 12 positions.

SDS-PAGE analysis of the SEC peak fractions confirmed the presence of a single band at the expected subunit molecular weight for each of the three successful designs (Fig. 2A, Table S1). Negative-stain transmission electron microscopy (NS-TEM) of the peak fractions revealed populations of well-dispersed, roughly spherical particles with diameters of approximately 150 Å (15 nm) for both designs, consistent with the dimensions predicted from the computational models (Fig. 2B). The particle images showed apparent 2-fold and 3-fold symmetry views characteristic of tetrahedral objects, supporting the intended cage architecture (Fig. 2C). No large aggregates or fibrillar structures were observed.

**Figure 2.**
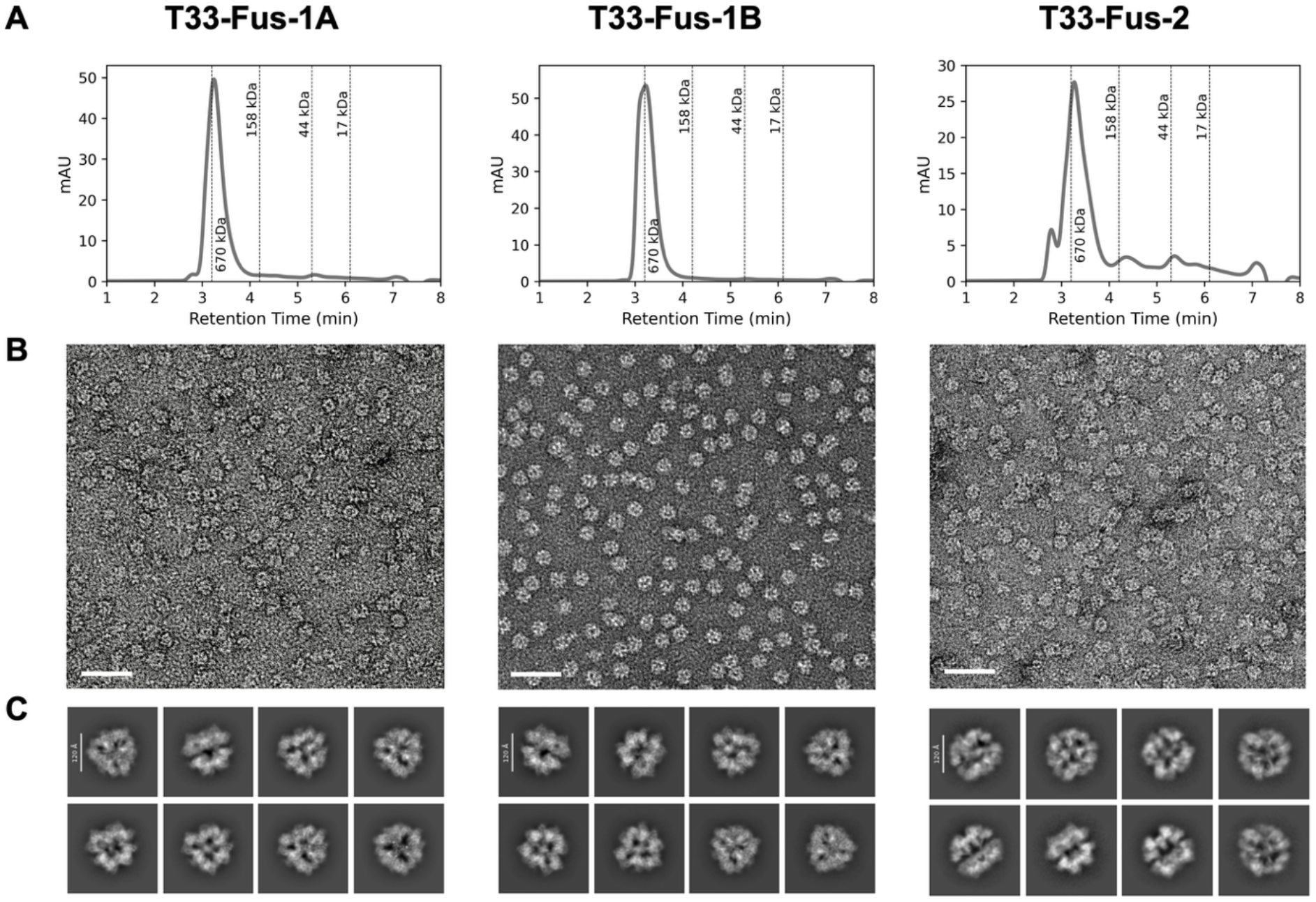
Experimental characterization of assembled protein cages. **(A)** SEC chromatograms of the three successful designs showing a single major peak at the expected elution volume for a 12-subunit assembly. **(B)** Representative negative-stain TEM micrographs revealing ~150 Å particles with apparent tetrahedral symmetry. Scale bar is 50 nm. **(C)** 2D classes showing different views of the particles from the cryo-EM data.

### Cryo-EM Structure Determination and Identification of Assembly Intermediates

All three SEC-positive designs (T33-Fus-1A, T33-Fus-1B and T33-Fus-2) were taken forward to high-resolution cryo-EM analysis. Purified proteins were vitrified on Quantifoil grids and imaged on a 200 kV Glacios microscope equipped with a Falcon 4 detector. Image processing and 3D reconstruction yielded density maps at overall resolutions of 3.0 Å (T33-Fus-1A), 3.9 Å (T33-Fus-1B) and 3.8 Å (T33-Fus-2), determined by the gold-standard FSC criterion at the 0.143 threshold (Fig. 3A, 3E, Fig. S1, S2). The refined atomic models (Fig. 3B, 3F) confirmed the intended tetrahedral T33 architecture. The maps showed clearly resolved secondary structure elements, side-chain densities for bulky residues, and the continuous α-helical linker connecting the two trimeric domains, enabling unambiguous model building (Fig. 3C, 3G). Because the structures of T33-Fus-1A and T33-Fus-1B are nearly identical at the backbone level while the resolution for the former is significantly better, further detailed analysis on this construct was based on T33-Fus-1A.

**Figure 3.**
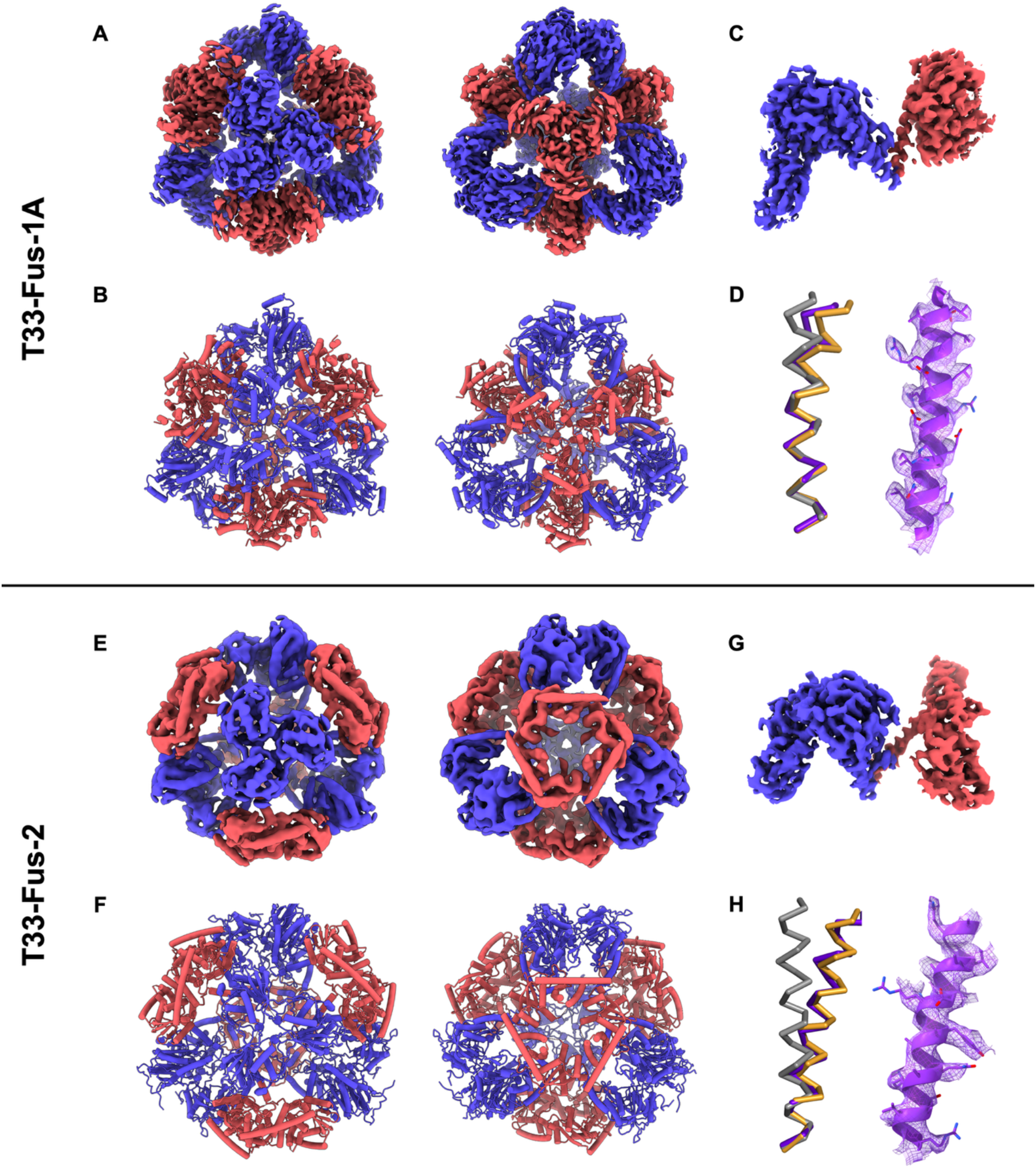
Cryo-EM structures of designs T33-Fus-1A and T33-Fus-2. **(A, E)** Cryo-EM density map viewed along the 3-fold axis of the N-terminal trimer domain (left) and 180° rotated cryo-EM map to show the 3-fold axis of the C-terminal trimer domain (right) for T33-Fus-1A and T33-Fus-2, respectively; **(B, F)** Refined atomic model showing the tetrahedral arrangement of two distinct trimeric domains (N-terminal trimer, blue; C-terminal trimer, red) connected by helical linkers, viewed along the 3-fold axis of the N-terminal trimer domain (left) and 180° rotated model to show the 3-fold axis of the C-terminal trimer domain (right) for T33-Fus-1A and T33-Fus-2, respectively; **(C, G)** Close up view of the cryo-EM structure for a single subunit for T33-Fus-1A and T33-Fus-2, respectively; **(D, H)** An alignment of the modeled alpha helical linker (orange) in the T33-Fus-1A (D) and T33-Fus-2 cage (H), the observed structure of the alpha helix (purple), and an idealized alpha helix (grey) (left). The helices were aligned by optimal overlap at their N-terminal five amino acids. The deviation at the C-terminal end highlights the helix bending that was introduced in the design model based on an algebraic treatment of alpha helix deformation (see Methods). The modeled bending was largely recapitulated in the observed cryo-EM structure. A fit of the refined structure to the density map is shown (right).

For T33-Fus-1A and T33-Fus-2, the experimentally determined structures agree remarkably well with the computational models. Superpositions of the cryo-EM structures onto the computational design models yielded overall backbone RMSD values of 0.92 Å and 2.51 Å for T33-Fus-1A and T33-Fus-2 respectively when overlapping a single subunit. Slightly higher RMS deviations are observed when overlapping the entire 12-subunit assemblies: 2.0 Å for T33-Fus-1A and 4.9 Å for T33-Fus-2. Given their diameters of approximately 170 Å in both cases, those deviations correspond to fractional errors of only 1.2% and 1.9% for the two constructs.

As expected, the agreement between observed and designed models is nearly exact for the trimeric cores, while relatively minor deviations occur in the helical linker region. The linker helix is continuous and well-ordered in the density maps for both constructs, indicating that the ProteinMPNN-designed junction sequences support stable helical geometry in the context of the assembled cage. Indeed, an examination of the helical fusion segments shows that the predicted helical bending, which was an important element in the design strategy, is recapitulated in the observed structures for both T33-Fus-1A (Fig. 3D) and T33-Fus-2 (Fig. 3H). The observed helical segments overlap much more closely with the slightly bent design models than they do with an idealized alpha helix. The T33-Fus-2 case is particularly striking; the deviation from ideal helix geometry is pronounced, while the match between model and observed geometry is remarkable (Fig. 3H).

## DISCUSSION

The present results demonstrate useful advances for applying the helical fusion strategy to protein cage design. The consideration of permissible modes of helix bending or deformation allows for more rational geometric scoring of candidate designs, while ML-based sequence design provides a rational approach for choosing amino acid sequences at positions where non-native residues are called for (e.g. at incidental contacts or in the linker itself where there is no native sequence). Prior structural studies on fusion-based cages provide important points of comparison for both technical advances introduced in the present study.

Concerning the issue of helical deformation, a series of structural studies on prior fusion-based assemblies emphasized the presence of irregularly shaped assemblies or alternative assembly forms. The first designed protein cage by Padilla *et al*. in 2001 is notable (14). A series of structural studies (38, 39) showed that diverse forms were typically obtained in X-ray crystallographic experiments. These were generally the result of compression and asymmetric deformation, sometimes reaching 18 Å RMS deviations. Studies on other fusion cages showed deformations sufficient to cause alternative assembly: symmetry T and D3 instead of O (17) and symmetry D3 instead of T (40, 41). In these prior cases of helical fusion (with the exception of Lai *et al*. 2015) (40), the natural oligomeric components appeared to be intact, with unexpected helical deformation being the cause of the design deficiencies. The structures obtained in the present study, where natural helical variation was analyzed during design, are much more robust to those deficiencies. The successful designs elucidated here are only a subset of the total designed sequences that were experimentally tested. Nonetheless, they represent the first fusion-based cages we are aware of where well-ordered, homogenous and symmetric architectures could be obtained in abundance and characterized in atomic detail.

Similarly, the importance of choosing optimal sequences in and around the helical fusion is made clear by the history of fusion-based cages. Polymorphism that challenged the structural characterization of the original design (14) was eventually overcome by judicious mutations in the linker region (37–39). Those studies predated robust software for sequence design. The present work, exploiting some of the latest machine learning tools for sequence design at key positions, led to highly ordered and symmetric assemblies.

Looking forward, the computational pipeline employed here is readily extensible. Higher-order symmetries (octahedral, icosahedral) and alternative oligomer combinations (e.g., trimer-tetramer or trimer-pentamer fusions) could be explored using the same framework. Integration with generative backbone design tools such as RFdiffusion (28) could further expand the accessible design space by enabling de novo generation of protein domains suitable for joining by helical junction backbones rather than relying solely on natural oligomeric components of known structure. Indeed, other recent studies have exploited helical fusions in pipelines using machine learning methods (33, 35). In the context of possible extensions to the present work, ML methods could be further exploited to simultaneously consider alpha helix bending and sequence design, and to include structure prediction metrics in scoring.

## CONCLUSIONS

Designed protein cages serve as platforms for a growing range of applications, including multivalent antigen display for vaccine development (1, 13), targeted drug delivery (3), protease-triggered disassembly for controlled release (42), and rigid scaffolds for cryo-EM imaging of small therapeutic proteins (43–45). Reliably engineering such novel architectures remains challenging. Improved strategies therefore offer expanded opportunities for successful development. Recent computational developments have played a critical role in expanding the set of available protein cages designed using interface design approaches (25–27), or de novo fold generation (28, 30, 46). The computational developments in the present work provide similar advances for fusion-based design approaches, in particular by incorporating ProteinMPNN into the sequence design stages (24). Notwithstanding the utility of ML methods demonstrated here and in other related work, classical mechanical principles remain important. As illustrated here, combining an algebraic or mechanical treatment of structural variability with modern ML methods diversifies the paths available for engineering robust protein cages and perhaps even more complex architectures.

The design and experimental work here provide two new protein cages (and sequence variations thereon), with homogenous and well-ordered three-dimensional structures not achievable before for fusion-based constructions. The protocol described, including with additional ML features, could be expanded to generate further sets of protein cages based on fusions. Besides not depending on the creation of novel protein-protein interfaces, fusion-based constructions offer additional counterpoints to interface-design methods. For example, the fusion approach typically involves single-component construction, whereas interface design approaches typically, though not always (see King *et al*. 2012) (21), involve multi-component design. In some settings, single subunit compositions are expected to offer manufacturing advantages. Fusion-type cages have also been noted to be particularly suited for conversion to protease sensitive forms with potential for triggered disruption (42). In addition, by avoiding the need for interface design, fusion-based approaches offer prospects for creating novel architectures where the complete sequence deviates from fully native sequences by only a small number of amino acids. That feature could be useful in future applications where limiting immunogenicity is a goal.

## METHODS

### Computational Design

#### Algebraic analysis of natural alpha helix deformation

Possible strategies for incorporating minor helix deformation into designed proteins include dynamical simulation, machine learning, clustering, and algebraic approximations based on known structural data. A strategy was developed here based on the latter. A set of alpha helical segments of length 7 (roughly 2 turns) were extracted from high resolution structures (dmin < 1.5 Å) in the PDB (36). That ensemble was then aligned spatially at the N-terminal three residues, and oriented in a canonical frame of reference based on the bond vectors from the first C-alpha to its amide nitrogen and its bonded carbonyl carbon. The pattern of observed deformation across the ensemble was then approximated as a problem of characterizing the rigid body space spanned by the set of i+6 residues from the ensemble. While rigid body space (in three dimensions) depends on six variables, for convenience the problem was treated in an augmented 9-dimensional space, based on the three x, y, z coordinates of three reference atoms (C-alpha, N and C atoms) from the i+6 residues. The symmetric 9×9 metric mass tensor from that ensemble was calculated, and the eigenvalues and eigenvectors extracted. A linear treatment of this type is suitable as long as rotational magnitudes are relatively small, as explained by Goldstein (47), and that was judged to be the case here. Accordingly, as expected, the three smallest eigenvalues (of 9) were small enough to be ignored (accounting for less than 0.2% of the variation), leaving 6 that embody the modes and magnitudes of natural helix variation, to the degree that that variation can be captured by the atomic positions at the ends of a semi-flexible segment. Interestingly, 90% of the total structural variation (i.e. the helix deformation) was captured by just the top 3 eigenvalues; 96% was captured by the top 4. The remaining two possible modes are practically unobserved. The finding that certain mathematically permissible modes of structural variation of the helix (e.g. direct stretching) are essentially forbidden is consistent with intuition and an understanding of atomic bonding but has not been described before in this form as far as we are aware. Having extracted the permissible modes and magnitudes of bending (in the form of atomic shifts of the i+6 residue in an established frame of reference), any helical conformation subject to some degree of non-ideality can be evaluated based on standard equations for probability using a quadratic Gaussian form. Note that this approach to characterizing bending was naive to underlying amino acid sequence effects and to the influences of surrounding structural features. The important interplay between those effects is recaptured in the sequence design step.

#### Construction of fusion models

Homotrimeric protein structures were retrieved from the PDB (36). Terminal α-helices were identified using DSSP (48). Combinatorial fusions were generated computationally by joining the helical ends of the two components, inserting an ideal helical linker with lengths up to 7 residues, or allowing the terminal ends of the two components to overlap by 2 residues (an effective linker length of −2). For each such potential construct, the symmetry axes carried by the two components were analyzed algebraically to obtain the angle between the axes and their separation at closest approach. Cases where those two values were within limits that might be mitigated by helix bending were retained for optimization. Perturbations to helix geometry were introduced according to the principal modes of variation obtained above. Specifically, the most probable deformation consistent with the exact symmetry requirements was obtained by sampling small deformation steps with magnitudes dictated by the eigenvalues associated with the principal modes and identifying the closest match to the symmetry requirements in each step. The total deformation was scored as a pseudo-energy based on the sum of squares of deformation magnitudes divided by normal mode eigenvalues. Candidate fusion subunits were evaluated for clashes between the two fused domains and between symmetry-related copies of the fusion subunit in the context of tetrahedral symmetry.

Sequences for the linker region and flanking positions were designed using ProteinMPNN (24). A total of 100 sequences were generated per backbone. Designs for experimental consideration were chosen by first filtering for the top 50% based on probability scores and then by scoring based on divergence from the consensus amino acid at each position. Designs closest to the consensus sequence overall were prioritized.

### Protein Expression and Purification

Genes encoding the designed fusion proteins were codon-optimized for *Escherichia coli* expression and synthesized by GenScript. Constructs were cloned into the pET-30a vector with a C-terminal His_6_ tag. Plasmids were transformed into *E. coli* LOBSTR-BL21(DE3) cells. Cultures were grown in TB medium supplemented with kanamycin at 37 °C to an OD_600_ of 1.2, induced with 0.25 mM IPTG, and incubated at 15 °C for 16 h. Cells were harvested by centrifugation, resuspended in lysis buffer (30 mM Tris, 200 mM NaCl, 1 mM TCEP, 20 mM imidazole, 0.1% LDAO,1 mM PMSF, pH 8.0) supplemented with benzonase nuclease, and lysed by sonication. Clarified lysate was applied to Ni-Charged MagBeads, washed with wash buffer (30 mM Tris, 200 mM NaCl, 1 mM TCEP, 20 mM imidazole, 50 mM sucrose, 0.01% Tween 20, pH 8.0), and eluted with elution buffer (30 mM Tris, 200 mM NaCl, 1 mM TCEP, 500 mM imidazole, 50 mM sucrose, 0.01% Tween 20, pH 8.0). The purified proteins were dialyzed into storage buffer (30 mM Tris, 200 mM NaCl, 5 mM EDTA, 1 mM TCEP, 50 mM sucrose, 0.01% Tween 20, pH 8.0) before being further purified by size-exclusion chromatography (SEC) on a Superose 6 Increase 10/300 GL column equilibrated in 30 mM Tris, 200 mM NaCl, 5 mM EDTA, pH 8.0.

### Negative-Stain Electron Microscopy

Purified protein at 0.05-0.2 mg/mL was applied to glow-discharged carbon-coated copper grids, blotted, and stained with 2% (w/v) uranyl acetate. Micrographs were collected on a Tecnai G2 Spirit microscope equipped with a 4kx4k TemCam-F416 CMOS camera (TVIPS) operating at 120 kV and on a 200 kV JEM-2200FS (JEOL) microscope equipped with a K3 direct electron detector (Gatan).

### Cryo-EM Sample Preparation, Data Collection and Processing

For cryo-EM, purified samples of designs T33-Fus-1A, T33-Fus-1B, and T33-Fus-2, were applied at a concentration of 0.3 mg/mL to glow-discharged Quantifoil R1.2/1.3 300 mesh copper grids, blotted for 3 s at 4 °C and 100% humidity using a Vitrobot Mark IV (Thermo Fisher Scientific), and plunge-frozen in liquid ethane. Data were collected on a Glacios microscope at 200 kV equipped with a Falcon 4. Movies were recorded at a calibrated pixel size of 0.93 Å/pixel with a total dose of ~40 e−/Å^2^.

All structures were determined using cryoSPARC (49). For T33-Fus-1A, a preliminary analysis in cryoSPARC Live was used to generate an *ab initio* model and a low-resolution refined map, which served as input for 2D template generation. A total of 4,200 movies were imported in EER format with 40 frames per movie. After patch motion correction and patch CTF estimation, micrographs were manually curated using thresholds of −0.5 to −3.5 µm average defocus and a maximum CTF-fit resolution of 4.5 Å, retaining 3,237 micrographs. Particles were picked with the template picker (200 Å particle diameter) using the 2D templates generated during cryoSPARC Live preprocessing and curated through particle inspection using local power and normalized cross-correlation thresholds. A total of ~413,000 particles were extracted in a 416-pixel box binned to 120 pixels. 2D classification was used to select ~109,000 particles, followed by two non-uniform refinement steps: the first in C1 (using the *ab initio* model from preprocessing as the initial volume) and the second with tetrahedral (T) symmetry imposed. The refined particles were re-extracted at lower binning (416-pixel box binned to 320 pixels) and subjected to several rounds of homogeneous and non-uniform refinement with T symmetry. The resulting volume served as input for per-particle reference-based beam-induced motion correction, and the particles were re-extracted unbinned in a 416-pixel box. These ~99,000 motion-corrected particles were subjected to further non-uniform refinement. To obtain a less anisotropic consensus map of the whole protein cage, the orientation of the particles was then rebalanced (retaining ~33,000 particles), yielding a 3.70 Å map. For focused refinement, the full (pre-rebalancing) particle set was subjected to symmetry expansion under T symmetry (~1.2 million expanded particles), locally refined, and then rebalanced (retaining ~435,000 particles). A final local refinement with a mask covering the asymmetric unit yielded a reconstruction at 2.99 Å.

For T33-Fus-1B, a similar workflow was followed. A total of 4,600 movies were imported into cryoSPARC in EER format with 40 frames per movie. A subset of 200 movies was preprocessed to generate an ab initio reconstruction and a preliminary low-resolution refined map for 2D template generation. After patch motion correction and patch CTF estimation of all movies, the micrographs were manually curated using thresholds of −0.5 to −3.5 µm average defocus and a maximum CTF-fit resolution of 4.5 Å, retaining 4,000 micrographs. Particles were picked with the template picker (200 Å particle diameter) using the templates generated during preprocessing. After particle inspection, ~451,000 particles were extracted in a 416-pixel box binned to 120 pixels. One round of 2D classification retained ~344,000 particles, which were used to generate an *ab initio* reconstruction. The best resulting map served as input for two non-uniform refinement steps: first in C1 and then with T symmetry imposed. The refined particles were re-extracted in a 416-pixel box binned to 320 pixels, and the ~295,000 retained particles were further refined by non-uniform refinement, followed by per-particle reference-based beam-induced motion correction. The orientation of the ~295,000 corrected particles was rebalanced (retaining ~184,000 particles) to obtain a consensus refined map by homogeneous refinement (4.24 Å). The particles were then symmetry-expanded under T symmetry (~2.6 million particles), and a final local refinement with a mask covering the asymmetric unit was performed, yielding a map at 3.91 Å resolution.

For T33-Fus-2, a subset of 1,200 movies was preprocessed to generate an *ab initio* reconstruction and a preliminary low-resolution refined map for 2D template generation. After patch motion correction and patch CTF estimation of all movies (4,086 movies), the micrographs were manually curated, retaining 2,893 micrographs. Particles were picked with the template picker (250 Å particle diameter) using the templates generated during preprocessing. After particle inspection, ~202,000 particles were extracted in a 320-pixel box binned to 96 pixels. One round of 2D classification retained ~47,000 particles, which were refined by homogeneous refinement, first in C1 and then with T symmetry imposed. The particles were re-extracted in a 320-pixel box binned to 256 pixels and refined again by homogeneous refinement with T symmetry, yielding a map at 5.11 Å. The particles were then symmetry-expanded under T symmetry (~561,000 particles), and a final local refinement with a mask covering the asymmetric unit was performed, yielding a map at 3.76 Å resolution.

Atomic models were built into the density using Coot (50) and refined with Phenix (51).

## Acknowledgements

We thank Duilio Cascio, Mark Arbing, Génesis Falcon, Alex Lisker, Michael Sawaya, Oscar Llorca, Rafael Fernández-Leiro, Lucas Tafur and the members of the CNIO Structural Biology Programme for valuable feedback and for providing helpful support. We thank Pablo Guerra from the IBMB-CSIC CryoEM Platform for assistance during the sample preparation and microscope data acquisition. We acknowledge funding from Project IU16-014045 (CRYO-TEM) from Generalitat de Catalunya and by “ERDF A way of making Europe”, by the European Union. We acknowledge ALBA Synchrotron for access and support of the cryo-EM facilities under BAG Proposal No CNIO-BAG 20250370388. We thank the Fundación Occident Visiting Researchers Programme at the CNIO. We thank Fundació Catalunya La Pedrera for its continuous support. R.C.-G. and P.S.-A. fellowships within the Generación D initiative, Red.es, Ministerio para la Transformación Digital y de la Función Pública, for talent attraction (C005/24-ED CV1), are funded by the European Union NextGenerationEU funds, through the Plan de Recuperación, Transformación y Resiliencia (PRTR). The Castells-Graells laboratory is supported by the Agencia Estatal de Investigación (AEI/10.13039/501100011033), Ministerio de Ciencia, Innovación y Universidades and co-funded by the European Regional Development Fund (ERDF-EU) [PID2024-161096NA-I00 to R.C.-G.], and is also supported by the Carlos III Health Institute (ISCIII) via CNIO.

