## Supplementary Material for "Design and structure of protein cages based on helical fusion and machine learning"

### Supplementary Figures:

a) T33-Fus-1A

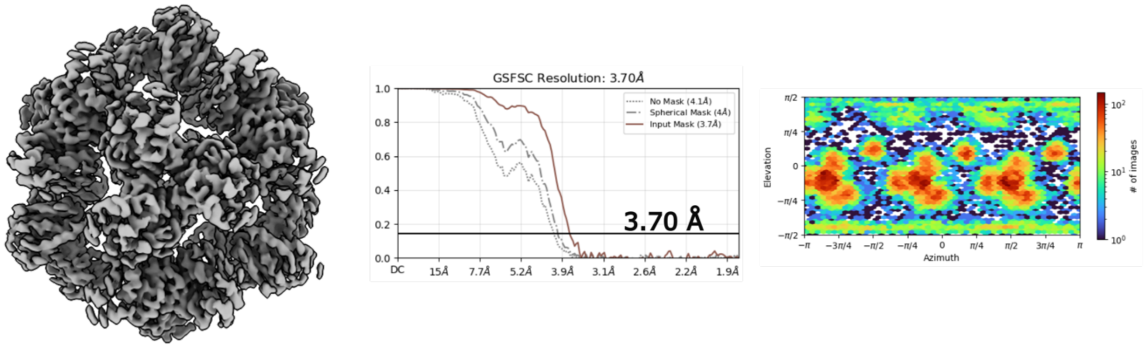


b) T33-Fus-1B

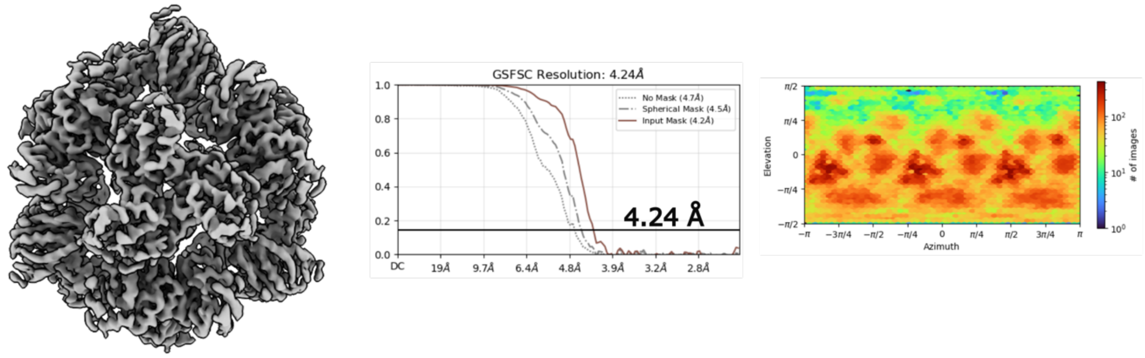


c) T33-Fus-2

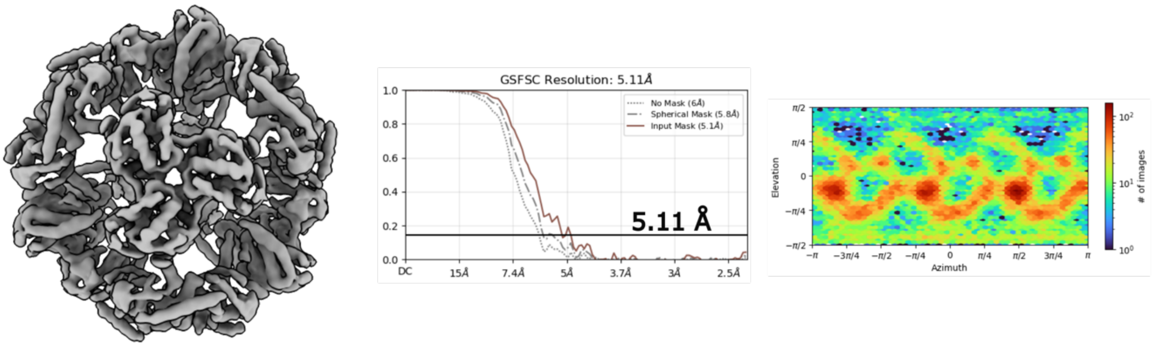


**Figure S1.** Summary of the cryo-EM maps with T symmetry. Cryo-EM consensus maps (T symmetry) reported in this manuscript with corresponding plots reporting the resolutions via gold standard FSC at a cutoff of 0.143 and the angular distribution of particles.

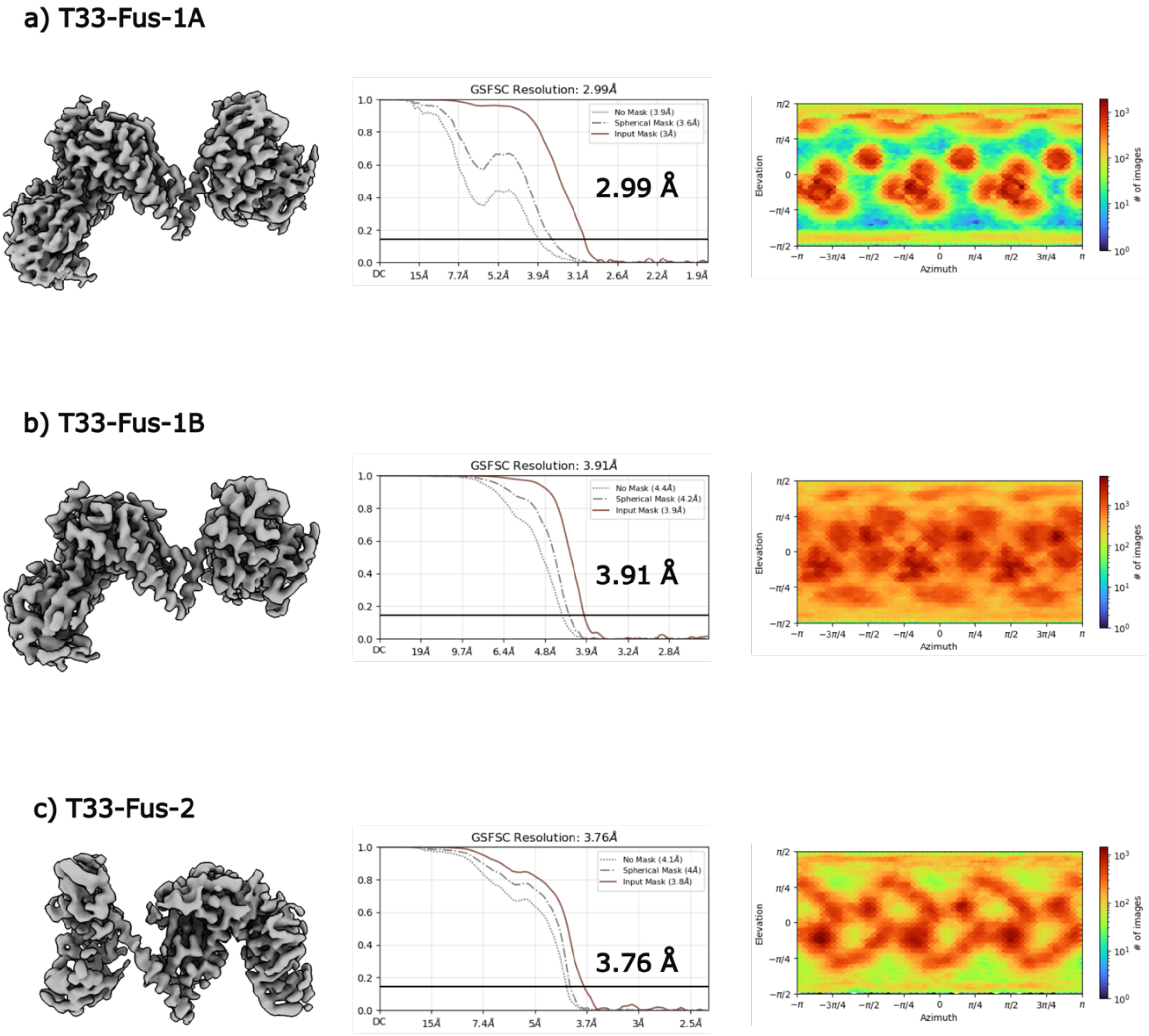


**Figure S2.** Summary of the cryo-EM maps of the asymmetric unit. Cryo-EM local refinement maps (C1 symmetry) reported in this manuscript with corresponding plots reporting the resolutions via gold standard FSC at a cutoff of 0.143 and the angular distribution of particles.

**Supplementary Text:**

<u>Protein sequences:</u>

| | |
|---|---|
| T33-Fus-1A | MDPQDLYTWEPKGLAVVDMALAQESAGLVMLYHFDGYIDAGETGD<br>QIVDQVLDSLPHQVVARFDHDRLVDYRARRPLLTFKRDTWSDYEEPT<br>IEVRLVQDATGAPFLFLSGPEPDVEWERFAAAVGQIVERLGVRLSVSF<br>HGIPMGVPHTRPVGITPHGSRTDLVPGHRSPFEEAQVPGSAEALVEY<br>RLAQAGHDVLGVAAHVPHYVARSAYPDAALTVLEAITAATGLVLPGIA<br>HSLRTDAHRTQTEIDRQIQEGDEELIALVRGLERRADGERIDAQLKAL<br>KVIPVIAIDNAEDIIPLGKVLAENGLPAAEITFRSDAAVEAIRLLRQAQPE<br>MLIGAGTILNGEQALAAKEAGATFVVSPGFNPNTVRACDEIGIPIVPGV<br>NNPSTVEAALEMGITTLKFFPAEASGGISMVKSLVGPYGDIRLMPTG<br>GITPSNIDNYLAIPQVLACGGTWMVDKKLVTNGEWDEIARLTREIVEQ<br>VNPGHHHHHH |
| T33-Fus-1B | MDPQDLYTWEPKGLAVVDMALAQESAGLVMLYHFDGYIDAGETGD<br>QIVDQVLDSLPHQVVARFDHDRLVDYRARRPLLTFKRDTWSDYEEPT<br>IEVRLVQDATGAPFLFLSGPEPDVEWERFAAAVGQIVERLGVRLSVSF<br>HGIPMGVPHTRPVGITPHGSRTDLVPGHRSPFEEAQVPGSAEALVEY<br>RLAQAGHDVLGVAAHVPHYVARSAYPDAALTVLEAITAATGLVLPGIA<br>HSLRTDAHRTQTEIDRQIQEGDEELIALVRGLEARYDGAVIDAQLKAL<br>KVIPVIAIDNAEDIIPLGKVLAENGLPAAEITFRSDAAVEAIRLLRQAQPE<br>MLIGAGTILNGEQALAAKEAGATFVVSPGFNPNTVRACREIGIPIVPGV<br>NNPSTVEAALEMGLTTLKFFPAEASGGISMVKSLVGPYGDIRLMPTG<br>GITPSNIDNYLAIPQVLACGGTWMVDKKLVTNGEWDEIARLTREIVEQ<br>VNPGHHHHHH |
| T33-Fus-2 | MREYEPGQPGMYELEFPAPQLSSSDGRGPVLVHALEGFSDAGHAIR<br>LAAAHLKAALDTELVASFAIDELLDYRSRRPLMTFKTDHFTHSDDPEL<br>SLYALRDSIGTPFLLLAGLEPDLKWERFITAVRLLAERLGVRQTIGLGT<br>VPMAVPHTRPITMTAHSNNRELISDFTPSISEIQVPGSASNLLEYRMA<br>QHGHEVVGFTVHVPHYLTQTDYPAAAQALLEQVAKTGSLQLPLAVLA<br>EAAAEVQAKIDEQVQASAEVAQVVAALERQYAAAQAKSAVARRLGK<br>VTASRVADVMTKTKSGYAASRQNYMAELIAQRLTGTQEIRFSNAAM<br>QRGTELEPHARARYIIETGEIVTEVGLIDHPTIAGFGASPDGLVGDTGLI<br>EIKCPNTWTHIETIKTGKPKPEYIKQMQTQMACTGRQWCDFVSYDD<br>RLPDDMQYFRTRIERDDALIAEIETEVSAFLAELEAEIEYLKRKAAKLA<br>GHHHHHH |

### Supplementary Table:

**Table S1.** Summary of the SEC purification values of the protein cages.

| Construct | SEC elution volume | SEC estimate MW | Subunit MW | Yield in mg (50 ml) |
|---|---|---|---|---|
| **T33-Fus-1A** | 3.2 min | 670kD | 52.82kD | 1.4 |
| **T33-Fus-1B** | 3.2 min | 670kD | 52.75kD | 2.86 |
| **T33-Fus-2** | 3.3 min | 640kD | 53.18kD | 0.496 |
